# Spatial transcriptomics uncovers hybrid, pro-inflammatory and pro-fibrotic cellular niches in pulmonary granuloma of patients with chronic sarcoidosis

**DOI:** 10.1101/2025.02.11.636632

**Authors:** Leonard Christian, Hande Yilmaz, Jannik Ruwisch, Leon Giercke, Benjamin Seeliger, Jan C. Kamp, Sirvan Bayraktar, Raphael Ewen, Theresa Graalmann, Jan Fuge, Mark Greer, Fabio Ius, Tobias Welte, Jens M. Hohlfeld, Marius M. Hoeper, Jens Gottlieb, Naftali Kaminski, Antje Prasse, Danny Jonigk, Yang Li, Christine Falk, Lavinia Neubert, Jonas C. Schupp

## Abstract

**Background:** Sarcoidosis is a disease of unknown etiology characterized by the formation of immune cell accumulation (granuloma) in the lung and other tissues. Chronic sarcoidosis may lead to pulmonary fibrosis.

**Aim:** To unravel cellular niches within pulmonary granuloma of chronic sarcoidosis patients using spatial transcriptomics.

**Methods:** Spatial transcriptomics using the Visium platform (10x Genomics) was performed on nine granuloma-containing lung explants from sarcoidosis patients. Validation of gene expression was performed through immunohistofluorescence protein staining and RNA *in situ* hybridization.

**Results:** Spatial gene expression covered 30,587 gene expression spots and 173 granulomas. A CD68^+^ macrophage niche was localized in the center of the granuloma, with a CD3^+^ T and CD20^+^ B cell niche in close proximity, surrounded by a COL3A1^+^ fibroblast niche. In the central granuloma macrophage niche, expression of the pro-fibrotic macrophage genes *SPP1*, *CHIT1* and *CHI3L1* was observed, genes whose expression has recently been described for macrophages in idiopathic pulmonary fibrosis. Additionally, pro-inflammatory macrophage genes were expressed in the central granuloma niche: macrophages appear armed for lysosomal degradation and ready for phagocytosis. Inner granuloma niches showed high responsiveness to interferon gamma (IFN-γ), expressing a multitude of IFN-γ-induced genes. High collagen and *CTHRC1* expression were observed in granuloma fibroblasts niches, characteristics of pro-fibrotic lung remodeling. Ligand-receptor analysis identified pro-inflammatory and pro-fibrotic interactions between granuloma niches.

**Conclusion:** Taken together, macrophages in the center of the sarcoidosis granuloma form an armed-and-ready, hybrid pro-inflammatory and pro-fibrotic niche, supporting granuloma persistence through continuous IFN-γ-stimulation and fibrotic remodeling conducted by fibrotic fibroblasts surrounding the granuloma.

## Introduction

Sarcoidosis is a multi-organ immune disease characterized by non-caseating granuloma and local inflammation. Lungs and intrathoracic lymph nodes are the most often affected organs (1). In many patients, these granulomas resolve without specific treatment, while in around 20% of cases, chronic sarcoidosis may lead to lung fibrosis or organ failure (2,3). As granulomas are the histological hallmark of sarcoidosis, their formation, structure, and significance have been studied over the last decades (4,5). However, the precise interactions on a cellular level and factors playing a role in granuloma persistence remain obscure.

The first step towards granuloma formation is carried out by sessile alveolar macrophages, reacting to a specific, yet unknown antigen in the lung (6). Then, monocytes are attracted to the inflammatory region, forming the core of the granuloma, where they acquire an epithelioid phenotype or fuse to multinucleated giant cells (7). The middle layer of the granuloma consists of recruited lymphocytes, including CD8^+^ T cells, T-helper (Th) Th1 cells, and Th17/Th17.1 cells, and B cells (8). In patients with chronic sarcoidosis, fibroblasts accumulate in the periphery of the granuloma, forming the outer granuloma layer. Although the formation of a fibroblast layer may be an important step of fibrotic tissue remodeling, which drastically worsens the prognosis for sarcoidosis patients, only little advancements were made towards characterizing these fibroblasts (9–11).

In this study, we employed spatial transcriptomics to explore the expression profiles, potential origins and interactions between cellular niches within the pulmonary granuloma of patients with chronic sarcoidosis, which may ultimately contribute to pulmonary fibrosis.

## Methods

Methods are detailed in the supplement and only briefly summarized here. This study was approved by the local Institutional Review Board (10142_BO_K_2022). Basic patient characteristics can be found in supplemental table E1.

### Processing of human lung tissue samples

Biobanked lung specimens of nine sarcoidosis patients, who underwent lung transplantation and had provided informed consent, were studied. Formalin-fixed and paraffin-embedded (FFPE) tissue blocks were sectioned at a microtome. Sections were stained with hematoxylin and eosin (H&E) to confirm the presence of granuloma. RNA quality was assessed using the RNeasy FFPE kit (Qiagen) for RNA isolation and DV200 measurement on a bioanalyzer.

### Spatial transcriptomics

Spatial transcriptomics and library generation were performed using the Visium platform from 10x Genomics according to manufacturer’s instructions. DNA library sequencing was carried out on the NovaSeq 6000 (Illumina) using associated reagents and standard procedure.

### Visium data acquisition, processing and analysis

Processing and visualization of spatial expression data were done with the ‘Seurat’ package in R software. Nine Visium samples passed the quality checks. Cluster of cells were identified through a clustering algorithm based on shared nearest neighbor (SNN) modularity optimization. Marker genes for each niche were generated with the Wilcoxon rank-sum test. FDR adjusted P values less than 0.05 were considered statistically significant in this study. Downstream analyses included gene set enrichment analysis and ligand-receptor analysis to explore niche-niche signaling networks.

### Validation

Multicolor immunofluorescence protein stains and RNA *in situ* hybridization (ISH) were performed for validation genes identified from the spatial gene expression analysis.

## Results

To characterize the cellular niches of the chronic pulmonary sarcoidosis granuloma, we performed spatial transcriptomics using the Visium platform (10x Genomics) on nine samples featuring granuloma of adequate number and RNA quality (Figure 1A and B, supplemental figure E1). In total, 30.587 tissue-covered gene expression spots were analyzed covering 173 granulomas. Cell type annotation based on the expression of distinct marker genes identified 13 niches, including four granuloma associated niches (Figure 1C-E).

**Figure 1:**
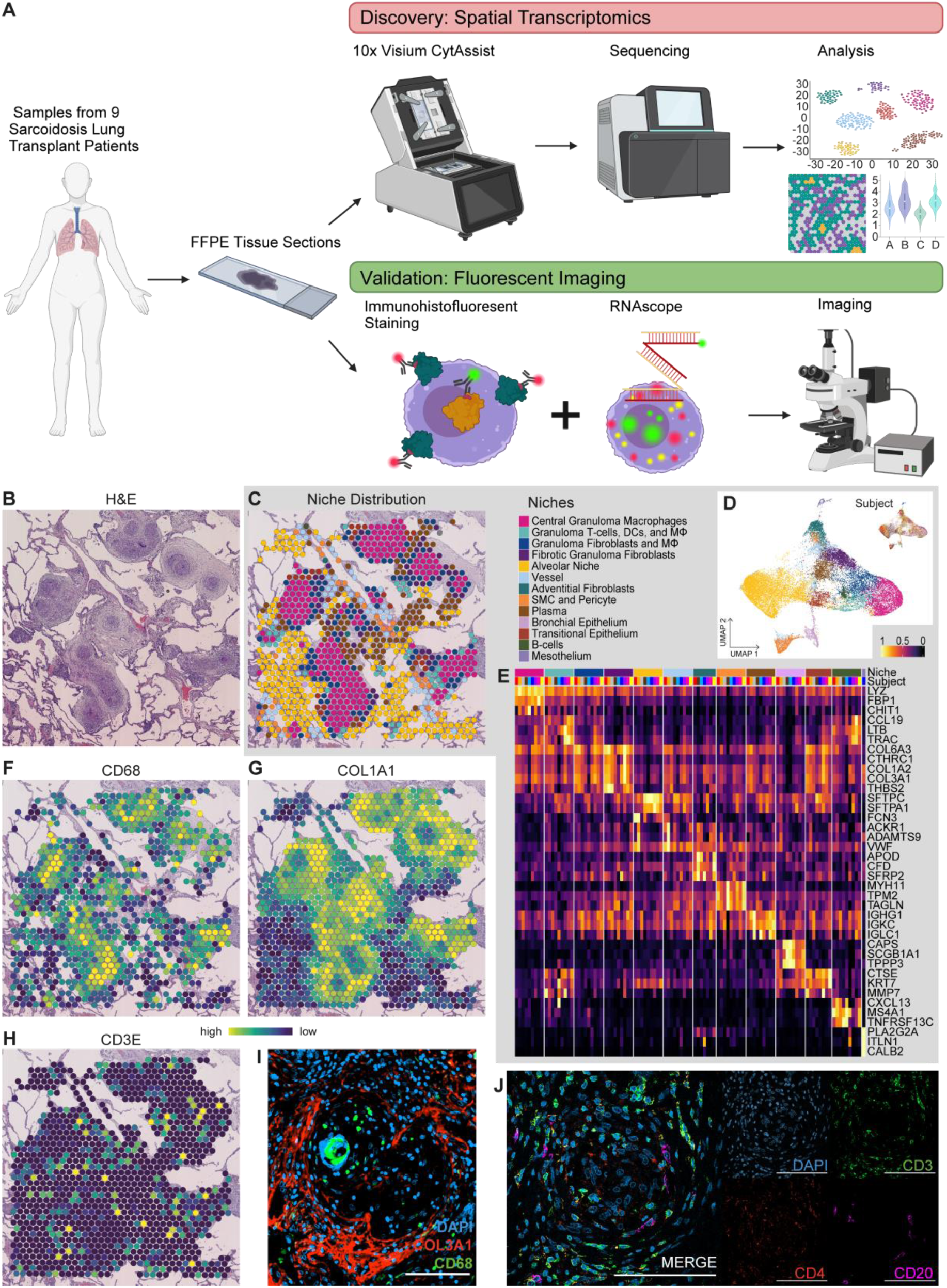
A) Biobanked FFPE tissue samples of nine chronic sarcoidosis patients undergoing lung transplantation were used. Spatial transcriptomics was performed on these samples using the Visium platform (10x Genomics). DNA libraries were sequenced and the data was analyzed. Based on spatial transcriptomics findings, validation via immunohistofluorescence protein staining (IHF) and RNA *in situ* hybridization (ISH) was performed. B) Exemplary Hematoxylin & Eosin (H&E) staining of a lung section containing granuloma. C) Spatial dimension plot showing the distribution of the cell type niches with each niche represented by a color-coded dot and niches identified based on marker gene expression in spatial transcriptomics. D) UMAP of spatial transcriptomics niches from all samples, with each niche color-coded and every spot representing a dot on a spatial plot. E) Heatmap displaying exemplary marker genes used to define the cell type niches in spatial transcriptomics. F) Spatial gene expression plot shows expression of macrophage marker *CD68* in the granuloma center. G) Spatial gene expression plot shows prominent expression of fibroblast marker *COL3A1* around the granuloma. H) Spatial gene expression plot shows weak expression of T cell marker *CD3E* around the granuloma. I) Representative immunohistofluorescence protein staining of macrophage marker CD68 (green) and fibroblast marker COL3A1 (red) and DAPI (blue) confirm the presence of macrophages in the granuloma and fibroblasts surrounding it (scale bar = 100 µm). J) Representative immunohistofluorescence protein staining of T cell markers CD3 (green) and CD4 (red), B cell marker CD20 (pink) and DAPI (blue) reveal the presence of T cells and B cells around the granuloma core (scale bar = 100 µm).

The main cell populations within the granuloma were uncovered based on marker gene expression, showing CD68^+^ macrophages in the granuloma center (Figure 1F) with a dense layer of COL3A1^+^ fibroblasts around it (Figure 1G). CD3^+^ T cells were loosely scattered in between the macrophage core and the fibroblast layer (Figure 1H). Low expression of dendritic cell marker genes (Supplemental figure E2) and B cell marker genes such as MS4A1 (CD20, figure 1E) were detected in predominantly in the T cell layer. The observed cellular distribution was confirmed through immunohistofluorescence protein staining (IHF) of CD68 and COL1A1 (Figure 1I) alongside staining for CD3, CD4 and CD20 (Figure 1J).

### Central granuloma macrophages occupy a hybrid, pro-inflammatory and pro-fibrotic niche

First, we explored the gene expression profile of granuloma macrophages, the main cells in the central granuloma niche. We observed high expression of *SPP1*, *CHIT1* and *CHI3L1* in the granuloma center (Figure 2A), genes associated with pro-fibrotic macrophages in idiopathic pulmonary fibrosis (IPF) (12), and markers of alternatively-activated (M2) macrophages (13,14). In parallel, high expression of *LYZ*, *IL1B*, *IL18* and *CYBB*, pro-inflammatory genes associated with conventionally-activated (M1) macrophages, was observed in the central granuloma niche (Figure 2B), suggesting that central granuloma macrophages exhibit both pro-fibrotic and pro-inflammatory characteristics. *GPNMB*, a novel marker for giant cells in sarcoidosis displaying characteristics of M1 and M2 macrophage polarization (15), was detected in the granuloma center (Figure 2A). Validation through IHF stains confirmed the presence of SPP1^+^ CHIT1^+^ macrophages in the central granuloma (Figure 2C). While SPP1 was distributed across the central granuloma, CHIT1 was colocalized specifically with CD68.

**Figure 2:**
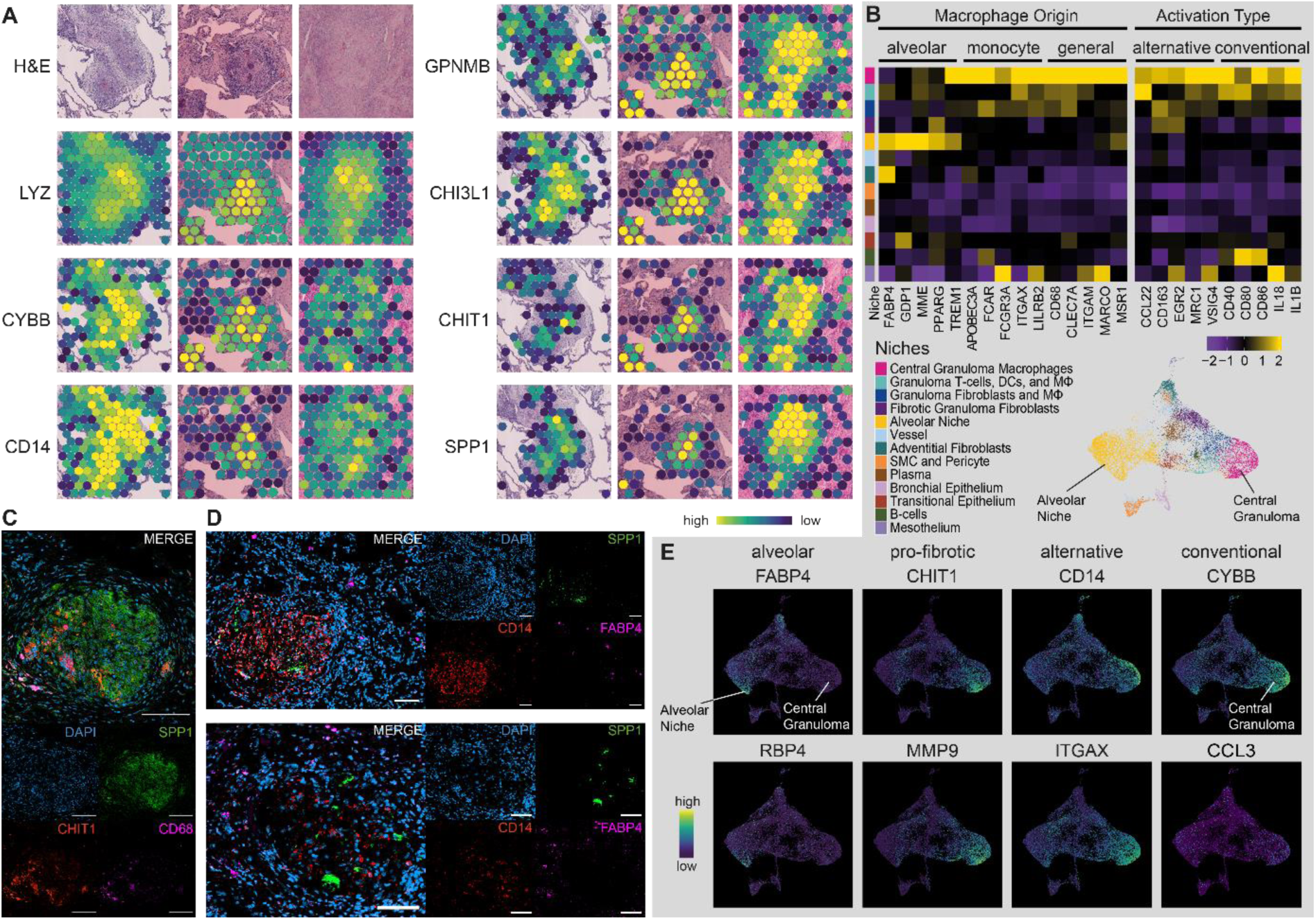
A) Spatial gene expression plot displaying genes associated with pro-inflammatory conventional macrophages (*LYZ* and *CYBB*), monocyte-derived macrophages (*CD14*) and pro-fibrotic alternatively-activated macrophages (*CHI3L1*, *GPNMB*, *CHIT1*, *SPP1*) in a granuloma. B) Heatmap featuring the gene expression across all subjects of pan-macrophage marker genes, macrophage origin marker genes (alveolar and monocyte), as well as markers for macrophage activation types alternative (M2) and conventional (M1). C) Representative immunohistofluorescence protein staining of the pro-fibrotic macrophage markers osteopontin (SPP1, green) and chitinase 1 (CHIT1, red), as well as macrophage marker CD68 (violet) and DAPI (blue) shows their presence in the center of the granuloma, but not in the surrounding tissue (scale bar = 100 µm). D) RNA *in situ* hybridization stains show the presence of *SPP1* (green) and *CD14* (red) in the center of the granuloma, while alveolar macrophage marker gene *FABP4* (violet) is expressed by tissue resident macrophages (scale bar = 100 µm). E) UMAP feature plot displaying the expression of macrophage marker genes across all niches. Alveolar macrophage marker genes *FABP4* and *RBP4* are highly expressed in the alveolar niche, while monocyte, alternative (M2) and conventional (M1) marker genes are expressed in the central granuloma niche.

To uncover the potential origin of granuloma macrophages, we examined the expression of alveolar and monocyte markers (Figure 2B and E)(16). While alveolar macrophage markers like *FABP4* were expressed in the alveolar niche, monocyte markers like *CD14* were expressed in the granuloma center. The expression of *FABP4* in alveolar, but not granuloma macrophages, and *CD14* in central granuloma macrophages alongside *SPP1*, was confirmed through ISH (Figure 2D).

Taken together, this data suggests that central granuloma macrophages are monocyte-derived, forming a hybrid, pro-inflammatory and pro-fibrotic niche.

### Central granuloma macrophages are primed to eliminate pathogens

To expand upon the inflammatory signaling within the central granuloma niche, we performed a pathway enrichment analysis using EnrichR, revealing associations with phagocytosis and endocytosis (GO:0050764, GO:0045807, and GO:0006896) as well as Toll-like receptor (TLR) signaling, phagosome, and lysosome pathways (Figure 3A and B). Expression of *TLR2* and its coreceptor *TLR1* was highest in the central granuloma niche (Figure 3C and F). Although *TLR4* expression was detected in the central granuloma, its expression was low compared to its coreceptors *LY96* and *CD14* and downstream TLR2/4 signaling genes, like *MYD88* and *IRAK1* (Figure 3C). Besides pathogen recognition, central granuloma macrophages displayed activity in pathogen uptake through expression of endocytosis-related genes such as *RAB5C*. Lysosomal degradation was especially prominent in the central granuloma, highlighted by expression of several lysosomal genes including *CTSB*, *CTSS*, *DNASE2*, *LIPA*, and *LYZ* (Figure 3C, D, and F). A shift to glycolysis and pentose phosphate pathway metabolism, facilitating quick immune cell proliferation and providing NADPH for oxidative stress response, was observed in granuloma macrophages through *HK3*, *GALM*, *TKT* and *FBP1* expression (Figure 3B and D), with confirmed protein expression of HK3 by CD14^+^ macrophages (Figure 3E).

**Figure 3:**
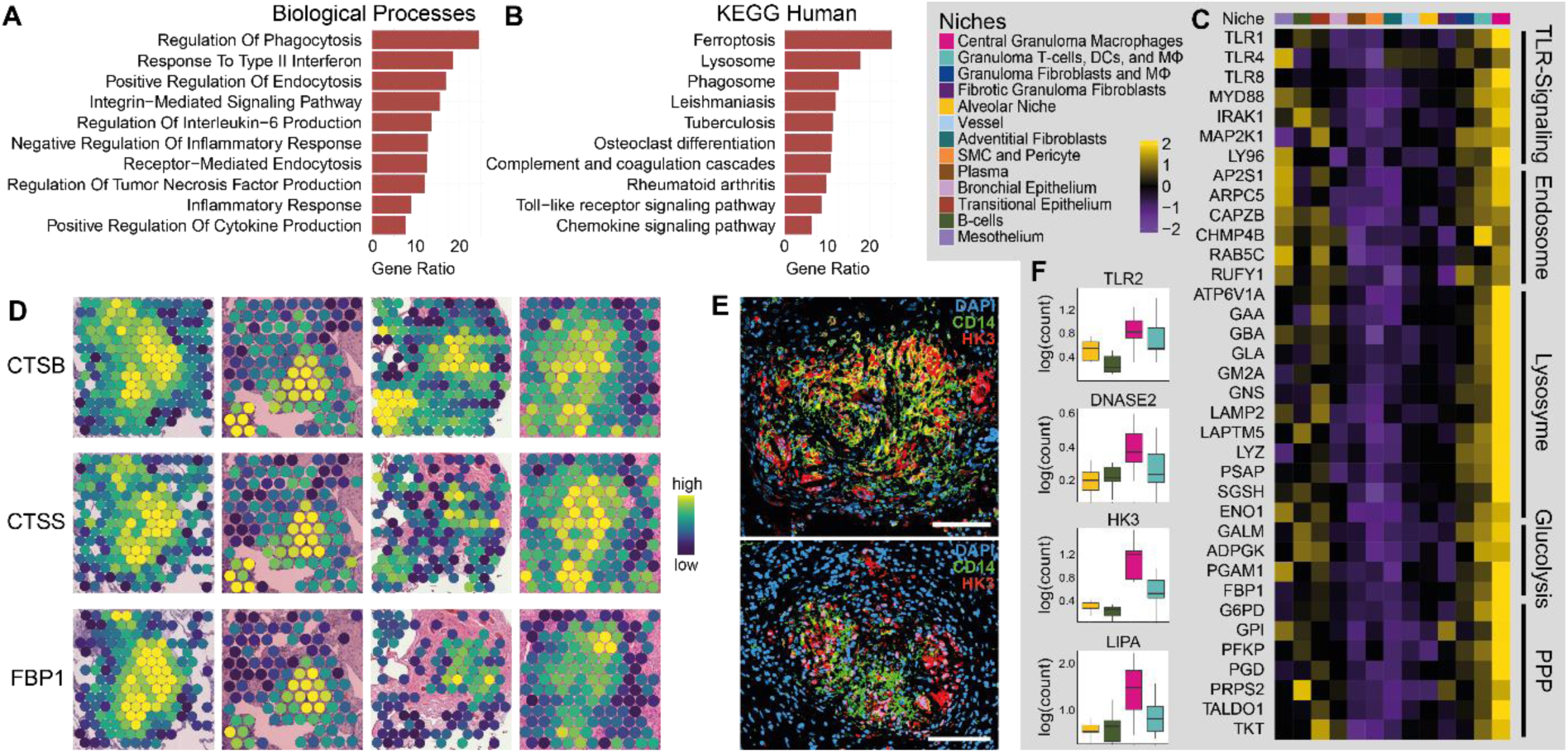
A) Gene ontology (GO) analysis of the top 200 genes expressed in the central granuloma niche by average log_2_(fold-change(FC)). Reference database: Biological Processes. B) Pathway analysis using the top 200 genes expressed in the central granuloma niche by log_2_(FC). Reference database: KEGG. C) Heatmap displaying the expression of genes involved in inflammatory pathways in the granuloma center. D) Spatial gene expression plot of lysosomal genes (*CTSB*, *CTSS*) and pentose phosphate pathway gene *FBP1* in a granuloma. E) Representative immunohistofluorescence protein staining of CD14 (green), HK3 (red), and DAPI (blue) shows that monocyte-derived macrophages in the center of the granuloma express glycolysis related genes (scale bar = 100 µm). F) Expression of genes involved in pathogen detection (*TLR2*), elimination (*LIPA*, *DNASE2*), and energy metabolism (*HK3*) is high in macrophage containing granuloma niches.

In summary, high expression of genes involved in pathogen recognition, uptake, and clearance alongside shifted energy metabolism suggests that macrophages in the central granuloma niche are primed to eliminate – potentially longe gone - pathogens.

### Interferon-γ signaling sustains a pro-inflammatory environment in the granuloma

Next, we examined how lymphocytes are involved in maintaining this pro-inflammatory and pro-fibrotic niche. Pathway analysis of the granuloma T cell, dendritic cell (DC) and macrophage (ΜΦ) niche revealed high responsiveness to IFN-γ by granuloma-associated cells (Figure 4A-C). These results are in line with high gene expression of IFN-γ-induced genes *CD74*, *CXCL9* and *IFI30* in the granuloma center (Figure 4D). Evaluating the expression of 50 genes induced by IFN-γ revealed specific spatial distribution of these genes across three different niches: i) central granuloma; ii) granuloma T cells, DCs, and ΜΦ; iii) B cell (Figure 4E). The IFN-γ-inducible chemokines *CCL5*, *CCL22* and *CXCL10* were prominently expressed in granuloma niches (Figure 4F), facilitating the recruitment of activated T cells and regulatory T cells (17,18). *CXCL9/CXCL10* and *CCL5/CCL22* expression were similar in the central granuloma and the granuloma T cells, DCs, and ΜΦ niche, underlining the ongoing lymphocyte recruitment to the granuloma. The highest expression of *CXCL16* and *IRF8* was observed in the central granuloma and the B cell niche, respectively. Taken together, pro-inflammatory IFN-γ signaling is maintained within pulmonary granuloma of chronic sarcoidosis patients.

**Figure 4:**
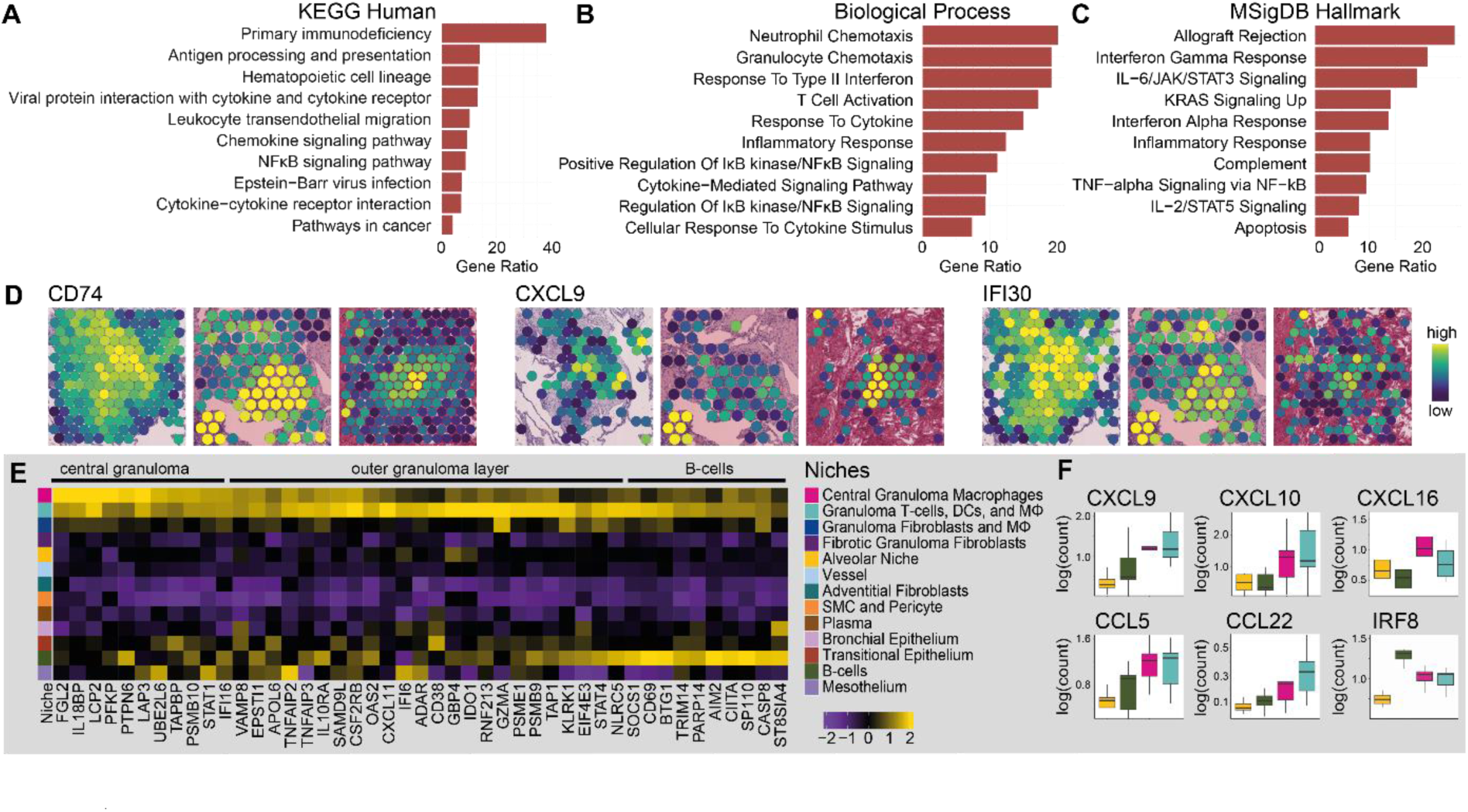
A) Gene ontology (GO) analysis of the top 200 genes expressed in the granuloma T cells, DCs, and ΜΦ granuloma niche by log_2_(FC). Reference database: KEGG. B) Gene ontology (GO) analysis of the top 200 genes expressed in the granuloma T cells, DCs, and ΜΦ granuloma niche by log_2_(FC). Reference database: Biological Processes. C) Pathway analysis of the top 200 genes expressed in the granuloma T cells, DCs, and ΜΦ granuloma niche by log_2_(FC). Reference database: MSigDB Hallmark. D) Spatial gene expression plot of interferon gamma (IFN-γ)-response-related genes in a granuloma. E) Heatmap displaying the expression of genes related to IFN-γ-response. F) Expression of genes involved in IFN-γ-response is high in granuloma macrophages, T cells, and DCs compared to surrounding tissue.

### The outer granuloma fibroblast niches contain CTHRC1^+^ fibrotic fibroblasts

Spatial transcriptomics revealed three fibroblast niches that were identified and classified by their location and gene expression: fibroblasts near small alveolar vessels (“adventitial fibroblasts”), fibroblasts in fibrotic lesions near granulomas (“fibrotic granuloma fibroblasts”), and fibroblasts around granulomas near immune cells (“granuloma fibroblasts and macrophages”).

Granuloma-associated fibroblasts showed no clear expression of adventitial and alveolar fibroblast markers, but expressed general fibroblast and myofibroblast markers (Figure 5A and C), while displaying prominent expression of the fibroblast marker genes *COL1A1*, *COL6A3*, *LUM*, and *SPARC* (Figure 5B). Fibrotic granuloma fibroblasts expressed high amounts of *CTHRC1*, which has recently been associated with IPF myofibroblasts (19,20). The granuloma fibroblasts and macrophages niche was characterized by its expression of *THY1* and *TNC* (Figure 5A). Additionally, all granuloma related niches showed high expression of *FAP*, a marker for collagen-producing fibroblasts in IPF (21,22). Pathway enrichment analysis of the fibrotic granuloma fibroblast niche revealed high levels of extracellular matrix (ECM) remodeling (GO:0030198, Figure 5D). ISH confirmed the expression of *COL1A1*, *CTHRC1*, and *TNC* in fibroblasts surrounding the sarcoid granuloma (Figure 5E). In summary, ECM remodeling was very active around the sarcoid granuloma and granuloma-associated fibroblasts show similarities to fibroblasts in IPF.

**Figure 5:**
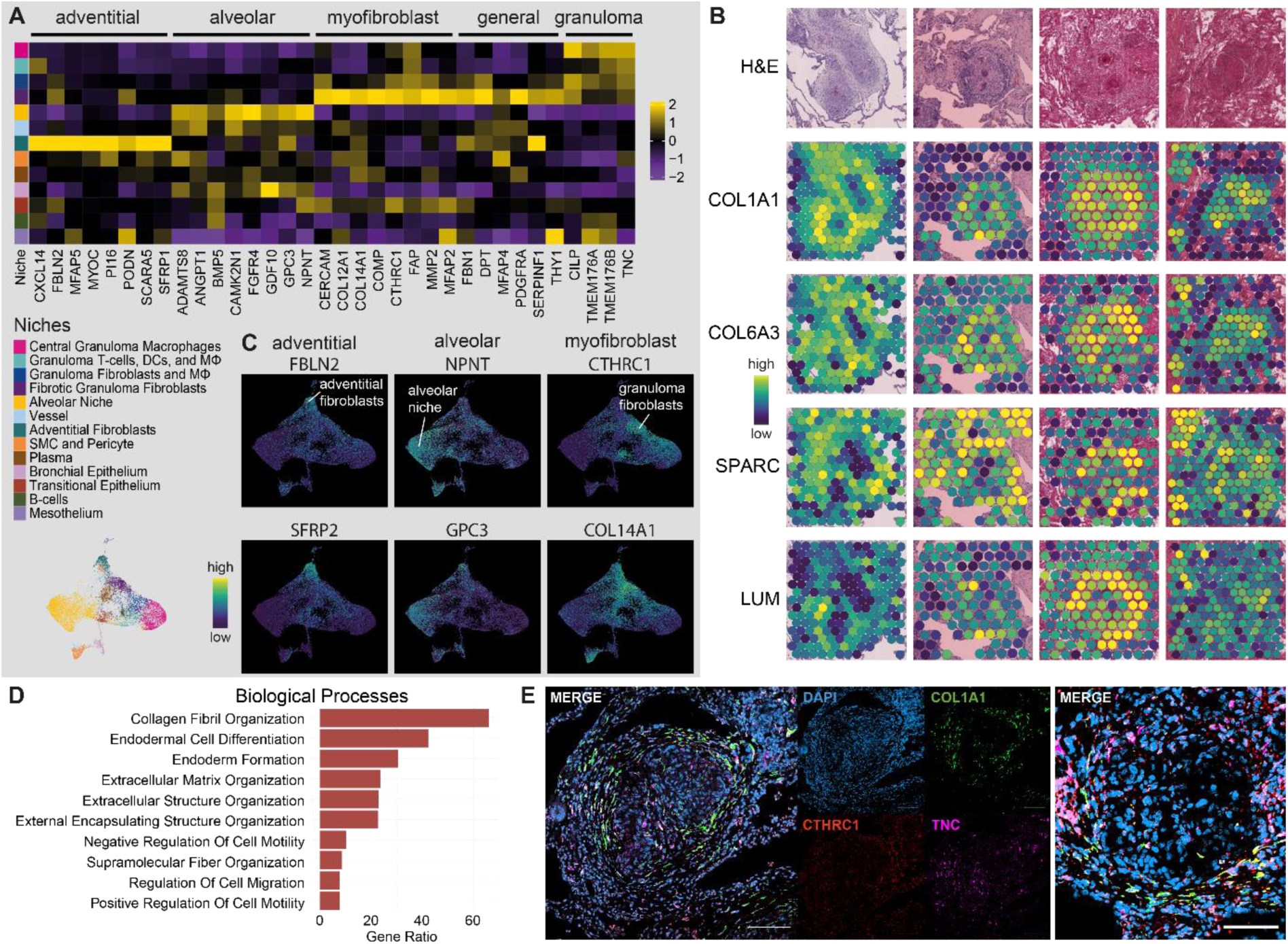
A) Heatmap displaying the expression of marker genes for adventitial fibroblasts, alveolar fibroblasts, and myofibroblasts, as well as pan-fibroblast markers and genes expressed primarily by granuloma fibroblasts and macrophages. B) Spatial gene expression plot of fibroblast-associated genes around a granuloma. C) UMAP feature plot displaying the expression of fibroblast marker genes across all niches. Adventitial fibroblast markers (*FBLN2*, *SFRP2*) are expressed in the adventitial fibroblast niche, alveolar fibroblast markers (*NPNT*, *GPC3*) are expressed in the alveolar niche, and fibrotic myofibroblast associated genes *CTHRC1* and *COL14A1* are expressed in the fibrotic granuloma fibroblast niche. D) Gene ontology (GO) analysis of the top 200 genes expressed in the fibrotic granuloma fibroblast niche by log_2_(FC). Reference database: Biological Processes. E) RNA *in situ* hybridization stains show the presence of *COL1A1* (green), *CTHRC1* (red), and *TNC* (violet) surrounding the granuloma. Nuclei are stained with DAPI (blue); scale bar = 100 µm.

### Pro-inflammatory and pro-fibrotic ligand-receptor interactions support granuloma integrity

Previous results demonstrated the expression and role of pro-inflammatory and pro-fibrotic genes in the granuloma, however, their importance in the crosstalk between granuloma niches and involvement in homeostasis and maintenance of the granuloma remained inconclusive. To bridge this gap, we aimed to further characterize this crosstalk employing ligand receptor analysis of the four granuloma associated niches distinguished in this study.

This approach confirmed pro-inflammatory signaling of the central granuloma niche via *CCL5* alongside pro-fibrotic signaling through *TGFB1*, *SPP1*, and *MMP9* (Figure 6A). *SPP1* and *MIF* signaling were found to be initiated by the central granuloma niche as the main sender, with interaction partners like *CD44*, *CD74*, and *CXCR4* being expressed in all granuloma niches (Figure 6B). Complement signaling via *C3* was sent by all niches, as highlighted by the strong expression of *C3* in and around the granuloma, with its receptors *ITGAX* and *ITGB2* being expressed predominantly in the central granuloma niche (Figure 6C). Signaling via CXC ligands showed strong involvement of *CXCL12*, whose receptor *CXCR4* was expressed in all granuloma-associated niches except the fibrotic fibroblast niche (Supplemental figure E5). Collagen and CXC ligand interactions were especially prominent in fibroblast containing outer niches, due to the strong gene expression of the associated ligands *COL1A1*, *COL1A2*, and *CXCL12* (Figure 6B). Spatial gene expression showed overlapping expression of ligand receptor pairs like *SPP1* and *CD44* within the boundaries of the granuloma (Figure 6C). Taken together, active pro-inflammatory and pro-fibrotic signaling persists within chronic sarcoid granuloma, with SPP1, CXCL chemokine, and collagen interactions playing a central role in this microenvironment.

**Figure 6:**
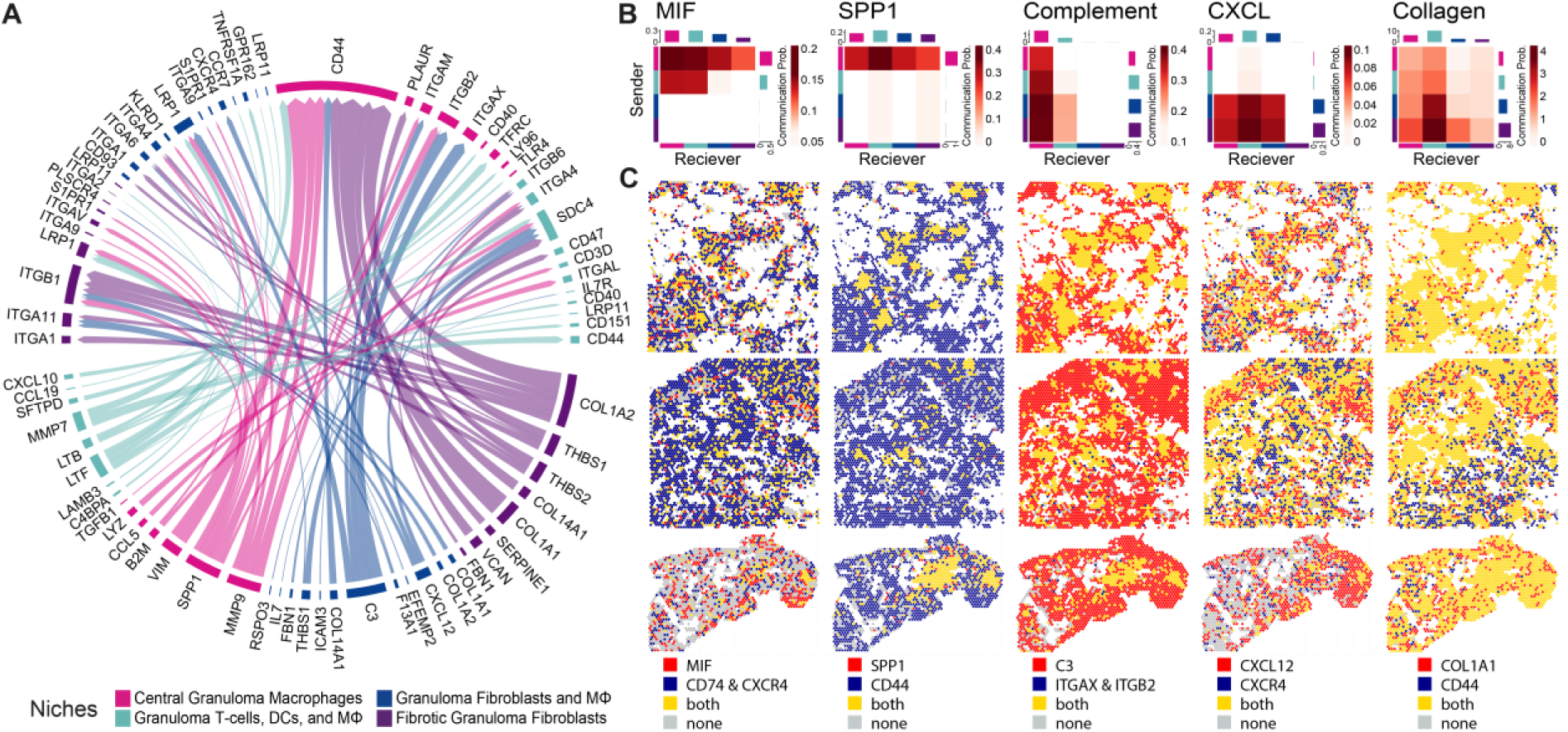
A) Circos plot showing the main ligand-receptor interactions within a granuloma. B) Heatmaps featuring ligand-receptor interactions within a granuloma. Depicted are pro-inflammatory (CXCL, Complement) and pro/fibrotic (SPP1, MIF, Collagen) signaling networks. C) Spatial gene expression of a ligand (red), its receptor (blue), the co-expression of ligand and receptor (gold) and neither ligand or receptor (gray).

## Discussion

In this study, we employed spatial transcriptomics on lung tissue samples from nine chronic pulmonary sarcoidosis patients to examine the gene expression of key niches in the granuloma, aiming to understand the mechanisms driving fibrotic remodeling and granuloma maintenance. Spatial transcriptomics revealed that macrophages in the central granuloma niche display hybrid characteristics of profibrotic and proinflammatory macrophages. These macrophages express pro-fibrotic genes like *SPP1*, *CHI3L1*, and *CHIT1*, while additionally expressing genes involved in pathogen detection and clearance, like TLRs, *LYZ*, and *LIPA*. Pathway analysis revealed that granuloma associated immune cells remain constantly stimulated by IFN-γ. Granulomas were surrounded by a prominent fibroblast layer, featuring high expression of collagens and *CTHRC1* with high ECM remodeling activity. Although the signal of T cells was weak compared to macrophages and fibroblasts, we were able to detect a distinct expression of receptors binding T cell derived chemokines in granuloma niches.

While the presence of pro-inflammatory and pro-fibrotic macrophages in the sarcoid granuloma have been observed before, their significance remains uncertain. A possible explanation is that a transition from pro-inflammatory to pro-fibrotic macrophages is concomitant with disease progression (23,24), as progressive, peri-granulomatous fibrosis is a typical characteristic of chronic sarcoidosis (3). Through this study, we found that genes associated with pro-inflammatory macrophages are still expressed within chronic sarcoid granuloma. In parallel, we observed that central granuloma macrophages resemble SPP1^+^ macrophages first described in IPF (12,19) and systemic sclerosis-associated interstitial lung disease (SSc-ILD)(25). SPP1^+^ macrophages in IPF similarly showed expression of *GPNMB* (26), alongside *CHIT1* and *CHI3L1*, which have been associated with development and progression of fibrosis in different kinds of interstitial lung disease (ILD) (27). High serum levels of chitinase 1 and YKL-40, encoded by the *CHIT1* and *CHI3L1* genes, respectively, have been shown to correlate with disease activity and progression in sarcoidosis (28–31). The expression of *SPP1*, *CHIT1 and MMP9* by granuloma macrophages was recently shown on a single cell level (32) and *SPP1*, *CHIT1* and *CHI3L1* were detected in the center of dermal granuloma from sarcoidosis patients (33). Ligand-receptor analysis demonstrated the expression of *SPP1* receptors involved in ECM remodeling in granuloma proximity, like *CD44*, supporting the role of *SPP1* in maintaining the granuloma structure.

The expression of pathogen recognition receptors by granuloma macrophages supports the hypothesis that stimulation of innate immune receptors triggers granuloma formation (8), while their expression in chronic sarcoidosis suggests a role in granuloma maintenance. Central granuloma macrophages expressed *TLR2*, which is involved in the detection of mycobacteria (34). Granuloma macrophages were not only able to detect mycobacteria, they also exerted high lysosomal activity, demonstrated by *LYZ*, *DNASE2*, and *LIPA* expression. Besides their function in lysosomal degradation, these genes are involved in granuloma formation through mTORc1/S6/STAT3 signaling (35–37). Additionally, we found expression of genes involved in glycolysis, a hallmark of disease progression in sarcoidosis (38), which also supports phagocytic and bactericidal activity management (39). Consistent with a recent report from a single cell sequencing sarcoidosis study (23), we observed high expression of pentose phosphate pathway-related genes in macrophages. As high pentose phosphate pathway activity is often observed during macrophage activation, this metabolic shift might promote granuloma formation in sarcoidosis. Whether the complement, MIF and CXC ligand interactions observed through ligand-receptor analysis might contribute to this pro-inflammatory environment remains unclear.

Despite the reported involvement of alveolar macrophages in the initiation of the granuloma formation (8), granuloma macrophages did not express alveolar macrophage marker genes. Instead, granuloma macrophages expressed monocyte marker *CD14*. This is in line with the findings that SPP1^+^ recruited macrophages were demonstrated to be derived from monocytes (40) and that monocytes are able to promote fibrosis in IPF (41). Considering the above, the dysregulated expression of pro-inflammatory genes in the central granuloma niche throughout granuloma maturation combined with pro-fibrotic genes might be the key to understanding the upkeeping of the granuloma.

Previous studies showed the importance of T cell-derived IFN-γ in sarcoidosis granuloma formation (42,43). While our assay did not detect IFN-γ itself, we found that *CXCL9*, *CXCL10*, and other genes involved in IFN-γ signaling pathways were expressed in both the central granuloma and the granuloma lymphocyte niches, indicating that IFN-γ signaling continues to play a role in sustaining established granulomas, supporting the potential use of drugs intercepting the IFN-γ pathway also in chronic patients. The importance of ongoing IFN-γ stimulation of these macrophages is underscored by evidence showing that elevated serum levels of *CXCL9* and *CXCL10* are linked to disease severity and help to regulate regulatory T cell recruitment (44,45) as well as to attract Th1 lymphocytes to the sarcoid lungs (46).

One of the most prominent complications associated with chronic sarcoidosis is loss of functional lung parenchyma due to fibrotic remodeling, which has been proposed to originate from the granuloma (47). Granuloma in patients with chronic pulmonary sarcoidosis were surrounded by fibroblasts, which displayed high expression of *CTHRC1*, a gene expressed in fibrotic lungs of IPF patients, but not in healthy people (20). CTHRC1^+^ fibroblasts have not been described for sarcoidosis yet, but may play a similar role in the development of fibrosis around pulmonary granuloma as their counterparts in IPF. *CTHRC1* has been proposed to be both an activator and inhibitor of Wnt/β-catenin signaling (15,48), suggesting a role in the differentiation of myofibroblasts participating in pulmonary fibrosis. This underlines the finding that granuloma-associated fibroblasts express myofibroblast associated genes, but only low amounts of alveolar or adventitial fibroblast associated genes. While *TNC* was described as an inducer of collagen expression in patients with SSc-ILD (49), is also an endogenous activator of TLR4 signaling (50), indicating that granuloma fibroblasts may harbor an undiscovered role in pro-inflammatory signaling within the granuloma.

This study is not without limitation. Statistically, our findings are mainly limited by the small sample size. Clinically, pulmonary manifestations are most common amongst sarcoidosis patients, granuloma forming in different organs, such as lymph nodes, skin, and heart, may differ from pulmonary ones. Additionally, we focused solely on chronic sarcoidosis cases that required lung transplantation due to disease severity. Technologically, the limited resolution of the Visium assay only allowed the characterization of cell type niches, not of single cells types, necessitating further in-depth studies on a larger scale.

Taken together, the sarcoid granuloma is a complex structure whose maintenance involves the crosstalk of diverse immune cells and fibroblasts. Using spatial transcriptomics, we explored the expression profile of the main niches within the chronic pulmonary sarcoid granuloma to elucidate their role within the granuloma. Our data suggests that macrophages present in the central granuloma niche possess seemingly contradictory functions, as they express genes involved in pathogen clearance and fibrotic ECM remodeling. While it is unclear if macrophages exhibiting a pro-inflammatory M1 phenotype later acquire the pro-fibrotic M2 phenotype or whether M2 macrophages accumulate in the central granuloma next to M1 macrophages forming an armed-and-ready M1/M2 hybrid niche, a fine balance of pro-inflammatory and pro-fibrotic factors appears to be the key of granuloma maintenance. Both SPP1^+^ macrophages and CTHRC1^+^ fibroblasts previously discovered in IPF emerged in our chronic sarcoidosis dataset. Translation from IPF research to sarcoidosis might alleviate chronic sarcoidosis research or open new avenues for sarcoidosis treatment in the future.

## Acknowledgments

The authors thank the Research Core Unit for Laser Microscopy and the Research Core Unit Genomics (RCUG) at Hannover Medical School for their support. The authors thank all study participants for their permission to use their respective tissue specimens for research.

