## Supplemental Data for "Spatial transcriptomics uncovers hybrid, pro-inflammatory and pro-fibrotic cellular niches in pulmonary granuloma of patients with chronic sarcoidosis"

### ONLINE DATA SUPPLEMENT

### SUPPLEMENTAL METHODS

#### Processing of human lung tissue samples

For this study, biobanked formalin-fixed and paraffin-embedded (FFPE) lung tissue blocks of nine sarcoidosis patients were used. Before explantation, patients provided informed consent for scientific research. These FFPE tissue blocks were cut on a routine microtome into 5 µm thick sections for spatial gene expression analysis, into 3 µm sections for immunohistofluorescence (IHF) staining and RNA *in situ* RNA hybridization (ISH), and into 50 µm sections for RNA quality control. Sections were positioned centrally on Superfrost Plus slides. Slides were stored dry at room temperature in a desiccator until use.

#### Sample assessment

Four 50 µm sections of each sample were collected in a tube for RNA quality assessment. RNA was extracted using the RNeasy FFPE kit (Qiagen, Cat. 73504) in accordance with the manufacturer's protocol. For this, tissue was rehydrated with deparaffinization solution (Qiagen, Cat. 19093). Tissue was lysed with Proteinase K (Qiagen, Cat. 73504) at 56°C for 15 min and then heated to 80°C for 15 min. Supernatant was treated with DNase I (Qiagen, Cat. 73504) for 10 min at RT. Buffer RBC (Qiagen, Cat. 73504) and 200 proof ethanol (Thermo Fisher Scientific, Cat. BP2818-500) were added to the lysate before applying the solution to an RNA-binding spin column. After washing with buffer RPE (Qiagen, Cat. 73504), the RNA was eluted in RNAse-free water. For RNA integrity evaluation, the DV200 value was determined at the Hannover Medical School Research Core Unit Genomics using a bioanalyzer high sensitivity-chip. In general, samples with DV200 values above 25% were considered for spatial transcriptomics.

Granuloma size and abundance were assessed trough hematoxylin and eosin (H&E) staining. FFPE sections subsequent to the spatial transcriptomics sections were rehydrated through a xylene and ethanol series. Hematoxylin (Sigma Aldrich, GHS116-500ML) staining was done with a 1:2 diluted Hematoxylin solution for 3 min. Slides were blued in tap water for 15 min. Sections were stained for 1 min in Eosin (Sigma-Aldrich, Cat. HT110116) and rinsed in ddH2O. Slides were then dehydrated and coverslipped. Samples containing distinct granuloma of adequate size and abundance were considered for spatial transcriptomics.

**Visium Spatial transcriptomics**

Spatial transcriptomics was performed with the Visium Spatial Gene Expression Reagent Kit (10x Genomics, Cat. 1000520) according to the manufacturer's instructions (CG000495 Visium CytAssist Gene Expression User Guide Rev A). In brief, sections were incubated at 60°C for 2h on a C1000 Touch™ Thermal Cycler equipped with the 96-deep well reaction module (Biorad, Cat. 1851197) prior to paraffin removal and tissue rehydration in a xylene (Carl Roth, Cat. CN80.1) and ethanol (Otto Fischar, Cat. 27690) series. Deparaffinized slides were stained with Hematoxylin (Sigma-Aldrich, Cat. GHS-1-16) for 3 min and bluing solution for 1 min (Fisher Scientific, Cat. 10381775) at RT. Slides were then stained with Eosin for 2 min at RT. In between these steps, slides were washed by dipping them in ddH20 according to the recommendations. Slides were coverslipped in 85% glycerol (Sigma-Aldrich, Cat. G9012-500ML) for tissue imaging. The coverslip was removed in ddH20 for destaining of the tissue by applying a 0.1 M HCl solution (Carl-Roth, Cat. T134.1) for 15 min at 42°C. Decrosslinking using buffer included in the kit (10x Genomics, Cat. 2000566) was performed for 1 h at 95°C.

Sections were incubated in PBS (Life Technologies, Cat. AM9624), 0.05% Tween-20 (Serva, Cat. 3979601) for 15 min before hybridization at RT. Left- and right-hand sides of the probes (10x Genomics, Cat. 2000657 and 2000658) in FFPE Hyb buffer (10x Genomics, Cat. 2000423) were added onto the tissue slides and incubated at 50°C overnight. After 20 h of incubation, slides were washed with pre-heated FFPE Post-Hyb Wash Buffer (10x Genomics, Cat. 2000424) thrice and in 2X SSC buffer (Ambion, Cat. AM9770) once. Probe ligation was performed for 1 h at 37°C with reagents provided by 10x genomics (10x Genomics, Cat. 2000425 and 2000445). Samples were washed with RT Post-Ligation Wash Buffer (10x Genomics, Cat. 2000419), incubated at 57°C, washed with buffer pre-heated to 57°C, and rinsed in 2X SSC buffer twice. Probe release was carried out on the Visium CytAssist instrument (10x Genomics, Cat. 1000441) using RNase (10x Genomics, Cat. 3000593) and Tissue Removal Enzyme (10x Genomics, Cat. 3000387). For CytAssist imaging, slides were stained with Eosin for 1 min. Probes were transferred from the tissue slides onto a Visium Slide (10x Genomics, Cat. 1000519) within the Visium CytAssist instrument. Slides were washed thrice with 2X SSC buffer. Probes extension was conducted for 15 min at 45°C using the extension enzyme and buffer provided in the kit (10x Genomics, Cat. 2000389 and 2000409). Slides were washed with 2X SSC buffer and released through 10 min incubation in 0.08 M KOH (Sigma-Aldrich, Cat. P4494-50ML). 1 M Tris-HCl (Life Technologies, Cat. AM9856) was added to each sample for KOH neutralization.

For pre-amplification of the probes, Amp Mix B (10x Genomics, Cat. 2000567) and TS Primer Mix B (10x Genomics, Cat. 3000511) were added to the samples. The pre-amplification PCR program was set to 98°C for 3 min, 10 cycles of denaturation at 98°C for 15 s, annealing at 63°C for 20 s, and extension at 72°C for 30 s with a final extension at 72°C for 1 min. PCR products were purified through size selection on AMPure beads (Beckman Coulter, Cat. A63881). Beads were washed with 80%

ethanol (Fisher Scientific, # 16606002). DNA was eluted with Buffer EB (Qiagen, Cat. 19086). To avoid over-amplification of the PCR product, a qPCR was performed to determine the DNA yield after the first amplification. SYBR FAST qPCR Master Mix (KAPA Biosystems, Cat. KK4600) was used along primers included in the 10x Genomics Visium kit (10x Genomics, Cat. 2000537). For this step, parts of the samples were diluted 1:5 in nuclease-free water according to recommendations. The Cq-value was determined for each sample through the following qPCR program: heating to 98°C for 3 min, followed by 30 cycles of 98°C for 5 s and 63°C for 30 s. For the sample index PCR, Amp Mix B and an individual dual index TS (10x Genomics, Cat. 1000251) were used. The sample index PCR program was set to 98°C for 3 min, a number of cycles equal to the rounded Cq-value of the sample plus 2 of: denaturation at 98°C for 15 s, annealing at 63°C, extension at 72°C for 30 s, with a final extension at 72°C for 1 min. PCR product was purified through double-sided size selection using AMPure beads. For quality control, samples were sent to Hannover Medical School Research Core Unit Genomics for a high-sensitivity run on an Agilent Bioanalzyer was performed for each sample. Samples were stored at -20°C until sequencing.

**Sequencing**

Sequencing libraries were prepared according to the manufacturer’s protocol. Libraries were handed over to Hannover Medical School Research Core Unit Genomics for sequencing using the NovaSeq 6000 platform and the designated NovaSeq 6000 SP Reagent Kit v1.5 with 100 cycles (Illumina, Cat. 20028316). The sequencing depth was estimated for each sample based on the Visium slide spots covered by tissue according to 10x Genomics recommendations.

**Immunofluorescence protein staining**

Slides were rehydrated through a xylene and ethanol series. Tissue was decrosslinked in 95°C water with 1 % Antigen Unmasking Solution (Vector Laboratories, Cat. H3301) for 20 min. Afterwards, the slides were cooled down to RT in 1X PBS (Chemsolute, Cat. 8418) for 20 min. Nonspecific antibody binding sites were blocked through incubation with 2.5% v/v normal donkey serum (Biozol, Cat. JIM-017-000-121) or 2.5% v/v normal horse serum (Vector Laboratories, Cat. S-2012) for 20 min depending on the species of the secondary antibody. Slides were then incubated for 1 h with each primary antibody and rinsed twice with PBS for 2 min between each incubation period. DyLight™ 488 Anti-mouse-IgG and DyLight™ 594 Anti-rabbit-IgG secondary antibodies both contained in the VectaFluor™ Duet Immunofluorescence Double Labeling Kit (Vector Laboratories, Cat. DK-8828) were applied to the slides for 1 h and subsequently washed in 1X PBS. For a three primary antibody panel, donkey anti-rabbit-Rhodamine Red X (Biozol, Cat. JIM-711-295-152), donkey anti-mouse AlexaFluor™ 647 (Life Technologies, Cat. A32787), and donkey anti-rat AlexaFluor™ 488 (Life Technologies, Cat. A21208) were used as secondary antibodies instead, followed by an additional 1X PBS wash step and an 1 h incubation with a pre-conjugated antibody. To reduce background autofluorescence, Vector TrueView Autofluorescence Quenching Kit (Vector Laboratories, Cat. SP-8400-15) was applied to each slide according to manufacturer's instructions. Slides were rinsed in 1X PBS before mounting with DAPI-containing Antifade Mounting Medium (Vector Laboratories, Cat. H-1800-10) and coverslipping. Slides were stored at 4°C in the dark.

**RNA in situ hybridization**

3 µm thick sections were incubated in an oven for 1 h at 60°C. Slides were rehydrated through 2x 5 min incubation in xylene (Carl Roth, Cat. CN80.1) and 2x 2 min incubation in 100% ethanol (Otto Fischar, Cat. 27690). Slides were dried in an oven for 5 min at

60°C. Deparaffinized slides were treated with Hydrogen Peroxide (ACD, Cat. 322000) for 10 min at RT. Hydrogen Peroxide was washed off in ddH2O by moving the slides up and down for ten times, which was repeated with fresh ddH2O. Slides were immersed in beakers containing 1x target retrieval buffer (ACD, Cat. 322000) pre-heated to 95°C for 30 min, then rinsed in ddH2O for 1 min twice, and transferred to 100% ethanol. Slides were dried for 5 min at 60°C. Tissue was encircled with a hydrophobic barrier pen (Vector Laboratory, Cat. H-4000). RNAscope Protease Plus (ACD, Cat. 322381) was applied on the tissue for 30 min at 40°C. Afterwards and in between all subsequent incubation periods, the slides were washed in 1x RNAscope Wash Buffer (ACD, Cat. 310091) twice for 2 min. RNAscope probes (Bio-Techne, Cat. 413331, 420771-C2, 311271-C3, 401891-C4) were mixed in the ratio 50:1:1:1 (C1:C2:C3:C4) and pre warmed to 40°C for 10 min. 100 µL of the probe mix was applied onto the slides for a 2 h incubation at 40°C. Slides were stored overnight in 5x SSC (Life Technologies, Cat. AM9770). RNAscope Multiplex FL v2 Amp 1, 2 and 3 were applied in order for incubation at 40°C for 30 min, 30 min and 15 min, respectively. Opal fluorophores (Akoya Biosciences, Cat. FP1487001KT, FP1488001KT, FP1495001KT, FP1497001KT) were reconstituted according to manufacturer's instructions before diluting them 1:1000 in TSA Buffer (ACD, Cat. 322809). The signal for each individual channel, starting with channel C1, was developed through incubation with RNAscope Multiplex FL v2 HRP-C1 (provided in ACD, Cat. 323110) for 15 min at 40°C, incubation with the Opal Fluorophore assigned to the channel for 30 min at 40°C, and incubation with RNAscope Multiplex FL v2 HRP-Blocker (provided in ACD, Cat. 323110) for 15 min at 40°C. This process was repeated with the respective reagents for fluorescence channel C2, C3, and C4. Before development of the Opal 520 fluorescence signal, TrueBlack Lipofuscin Autofluorescence Quenching Solution (Biotium, Cat. B-23007) diluted 19:1 in 70% Ethanol was applied for 1 min.

DAPI (provided in ACD, Cat. 323110) was added to the slides before coverslipping with Prolong Gold Antifade Mounting Medium (Thermo Fisher Scientific, Cat. P36930). Slides were stored at 4°C in the dark until scanning.

**Immunofluorescence Microscopy**

Slides were scanned using the Zeiss Axio Scan 7 equipped with a 20x Plan-Apochromat 20x/0.8 27M objective. Emission was filtered using the 96 HE BFP 450/40, 38 eGFP 525/50, 43 HE DsRed 605/70, 26 Alexa Fluor 660 685/50, and 50 Cy 5 690/50 broad pass filters. For detailed images, the confocal laser scanning microscope LSM 980 Airyscan 2 (Zeiss), equipped with 405 nm, 488 nm, 514 nm, 561 nm, and 633 nm lasers, alongside a 40x oil immersion objective and two confocal PMT detectors operated by the Hannover Medical School Research Core Unit for Laser Microscopy was used.

**Visium data acquisition and processing**

Processing and visualization of spatial expression data were done with the 'Seurat' package (v.4.3.0.1) in the software R (v.4.2.3). We processed ten Visium H&E samples, of which nine passed quality standards. The Read10X_Image function was used to load images of the Visium samples, and the Visium Spatial Experiment data was imported into Seurat using the Load10X_Spatial function. Normalization of 30,587 raw counts was conducted with the NormalizeData function, setting the normalization method to "LogNormalize" and the scale factor to 10000. The nine Visium samples were then merged into a single Seurat object. Feature selection was performed to identify features with significant cell-to-cell variation using the FindVariableFeatures function with default parameters. Next, gene expression data was scaled using the ScaleData function. Principal component analysis (PCA) was conducted with the

RunPCA function to reduce the dataset's dimensionality. During clustering analysis, The FindNeighbors() and FindClusters() functions were applied sequentially, with the resolution set to 1. Cluster of cells were identified through a clustering algorithm based on shared nearest neighbor (SNN) modularity optimization. Non-linear dimensionality reduction was performed using the RunUMAP function.

**Identification and Visualization of Marker Genes**

Marker genes for each niche were detected with the Wilcoxon rank-sum test. Adjusted P values less than 0.05 were considered statistically significant in this study. To visualize the gene expression of marker genes as $\log_2$(FC) for every niche, a unity normalized average per subject heatmap was created with the function 'Heatmap' in the 'ComplexHeatmap' package (v.2.14.0). The gene expression across the niches was visualized with the function 'FeaturePlot' in the 'Seurat' package. A box plot was used to visualize gene expression across the niches using the function 'geom_boxplot' in the 'ggplot2' package (v.3.4.3). The 'coord_cartesian' range (ylim) was set to 0.1 to 2.0 to adjust the visible range.

**Expression of genes and the determination of spatial features**

We visualized spatially resolved gene expression for selected regions of the Visium dataset with the function 'SpatialDimPlot' and 'SpatialFeaturePlot' in the 'Seurat' package (v.4.3.0.1), with default parameters, adjusting the min. cutoff to "quantile 1" and max. cutoff to "quantile 95" to display gene expression variation across different regions of the tissue sample.

**Gene set enrichment analysis**

For Gene Set Enrichment Analysis the ‘ggplot’ function from the ‘ggplot2’ package (v.3.4.3) was employed, selecting the 200 genes with the highest counts by $log_2$(FC) of a certain niche as input. Gene symbols of these genes were converted to ENSEMBL IDs with the function ‘bitr’ from the package ‘ClusterProfiler' (v.4.8.3) and genome annotation for human org. ‘Hs.eg.db’ was used. Gene Ontology Biological Process (GO_Biological_Process_2023), Kyoto Encyclopedia of Genes and Genomes (KEGG_2021_Human), and The Molecular Signatures Database (MSigDB_Hallmark_2020) were used as reference datasets using the ‘enrichR’ function from the ‘enrichR’ package (version v.3.2). Only the ten significant gene terms with the highest gene ratio were displayed.

**Exploring ligand receptor signaling**

A combined set of unique ligand and receptor gene symbols derived from the ncomms8866_human dataset in the Connectome package were identified. These genes have been filtered to include only genes whose RNA is detectable via probes in Visium. Those genes were scaled with the function ‘ScaleData’ from the ‘Seurat’ package with the default parameters. A connectome was generated with the function ‘CreateConnectome’ from the ‘Connectome’ package (v.1.0.1). The data was filtered using the function ‘FilterConnectome’ from the Connectome package to identify edges with a ligand and receptor z-score > 0 and with both the ligand and receptor expressed in at least 10% of the cells in their respective niches. The data was grouped by the ligand receptor niches and then the Top 5 genes by weight scales were selected. The Circos plot was created using the ‘CircusPlot’ function from the ‘Connectome’ package, employing edge weights from the normalized slot and data were mapped against the FANTOM5 database.

To identify and analyze cell-cell communication networks, a CellChat object was created using the function ‘createCellChat’ in the ‘CellChat’ package (v.2.1.2), with a Seurat object as the input data.  The default CellChatDB.human reference database for analyzing ligand receptor interactions in human samples was used. To achieve this, the functions ‘subsetData’, ‘identifyOverExpressedGenes’, and ‘identifyOverExpressedInteractions’ were used with their default parameters. Communication probability was computed using the ‘computeCommunProb’ function, with the type set to 'triMean' (the default parameter) to produce fewer but stronger interactions. While the communication probability was computed at the signaling pathway level using the ‘computeCommunProbPathway’ function, the aggregated cell-cell communication network was calculated with the function ‘aggregateNet’.  Cell-cell communication network of MIF, SPP1, Complement, CXCL, and Collagen interactions were visualized using a heatmap with the ‘netVisual_heatmap’ function. Signaling genes associated with MIF, SPP1, Complement, CXCL, and collagen signaling pathways were identified using the ‘extractEnrichedLR’ function, and all significant interactions were visualized with a bubble plot using the ‘netVisual_bubble’ function. Gene expression distribution in granulomas was visualized with the function ‘spatialFeaturePlot’ in the ‘CellChat’ package.

The 'RunNICHES' function from the NICHES package (v.1.0.0) was applied to analyze cell-cell signaling in the data, using the Seurat object as input, along with the FANTOM5 database. The 'ScaleData' function was used to scale features, while the 'FindVariableFeatures' function from the Seurat package was employed to identify variable features. The 'selection.method' parameter was set to 'disp' to select genes with the highest dispersion values. Differentially expressed genes were analyzed using the 'FindAllMarkers' function from the Seurat package, with a $\log_2$(FC) threshold of 0.1

and the 'wilcox' test applied. The 'DoHeatmap' function in the Seurat package was used to generate a heatmap.

### SUPPLEMENTAL TABLES

**Table E1. Patient characteristics**. Basic characteristics of the sarcoidosis patients included in the dataset. Continuous variables are depicted as median and interquartile range. Age is given in years. FVC and DLCO are calculated as percent of the baseline values. All lung function values show last available lung function pre lung transplantation.

| **n** | **Sarcoidosis n = 9** |
|---|---|
| Female /Male [n] | 3/6 |
| Median Age at diagnosis in years [Q1/Q3] | 38 [33/40] |
| Median Age at lung transplant in years [Q1/Q3] | 56 [49-59] |
| Median †FVC % predicted [Q1/Q3] | 44 [39-55] |
| Median ‡DLCO % predicted [Q1/Q3] | 32 [19 - 35] * |
| Pulmonary Hypertension [yes/no] | 6/3 |
| Prednisolone [yes/no] | 4/5 |
| Tadalafil [yes/no] | 4/5 |
| Endothelin Blocker [yes/no] | 1/8 |

†DLCO=Diffusing capacity for carbon monoxide; ‡FVC= Forced vital capacity; *=missing values.

**Table E2. Primary antibodies used for immunohistofluorescence protein staining.**

| Target | Host | Distributor | Catalog | Dilution | Clone |
|---|---|---|---|---|---|
| CD3 | Rat | Bio-Rad | MCA1477T | 1:50 | CD3-12 |
| CD4 | Rabbit | Roche | 790-4423 | Ready to use | SP35 |
| CD14 | Mouse | Santa Cruz | sc-58951 | 1:50 | 5A3B11B5 |
| CD20 | Mouse | Bio-Trend | IHC532-100 | 1:100 | IHC532 |
| CD68 | Mouse | Santa Cruz | sc-20060 AF488 | 1:250 | KP1 |
| CD68 | Mouse | Life-Technologies | 14-0688-82 | 1:100 | KP1 |
| CHIT1 | Rabbit | Sigma Aldrich | HPA010575-25UL | 1:50 | Poly |
| COL3A1 | Rabbit | Proteintech | 22734-1-AP | 1:50 | Poly |
| HK3 | Rabbit | Sigma Aldrich | HPA056743-25UL | 1:50 | Poly |
| SPP1 | Mouse | Invitrogen | MA5-17180 | 1:25 | 7C5H12 |

**SUPPLEMENTAL FIGURES**

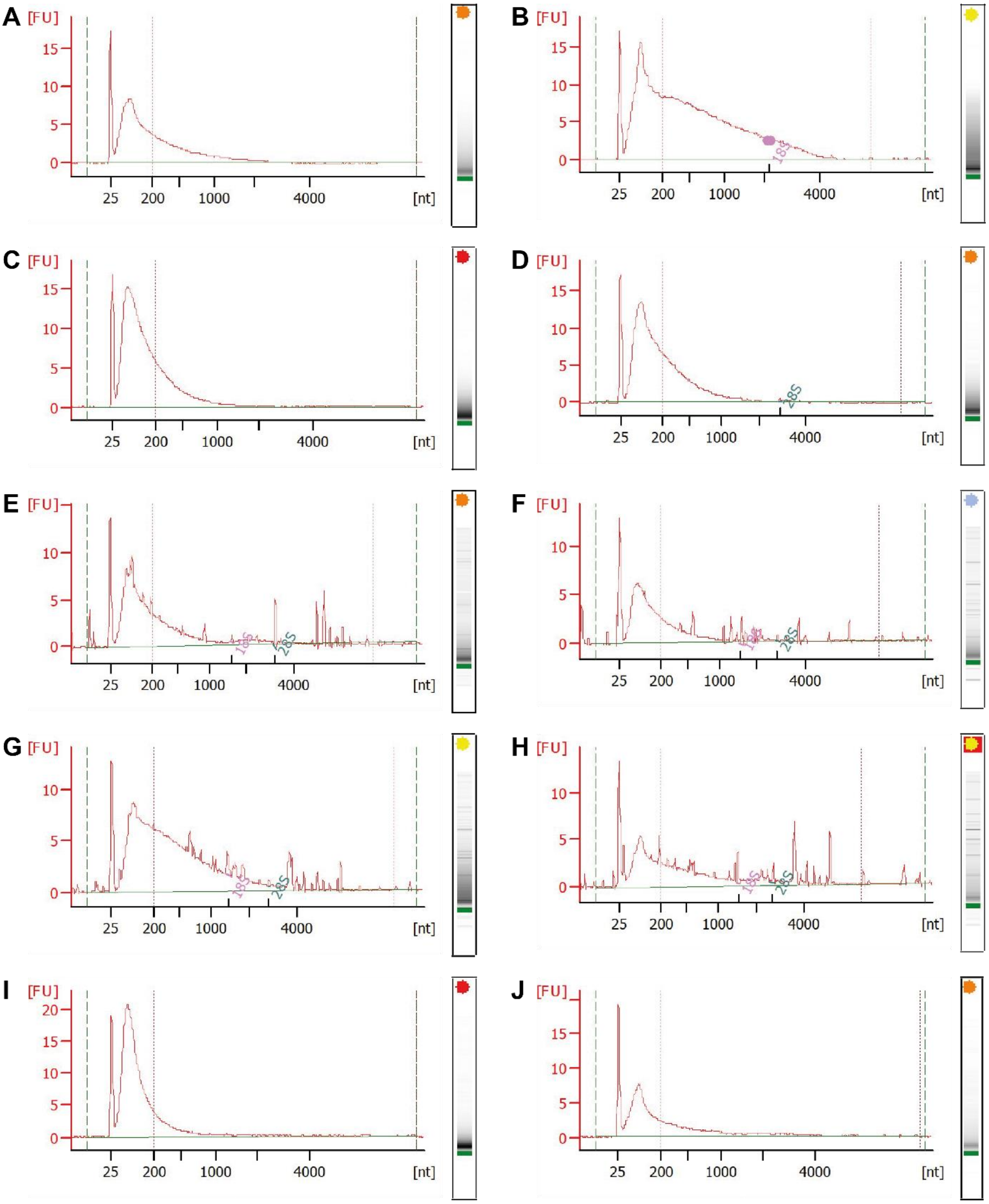


**Figure E1:** Fragment length measurement and quality assessment of RNA isolated from long-term stored FFPE blocks of sarcoidosis patients.

A) DV200 = 34%, RIN: 2.4, time since explantation: 4 years;

B) DV200 = 59%, RIN: 2.3, time since explantation: 2 years 15;

C) DV200 = 25%, RIN: 2.5, time since explantation: 9 years

D) DV200 = 34%, RIN: 2.4, time since explantation: 6 years

E) DV200 = 36%, RIN: 3.20, time since explantation: 0.5 years

F) DV200 = 35%, RIN: 5.60, time since explantation: 1 years

G) DV200 = 58%, RIN: 2.50, time since explantation: 3 years; This sample was removed from the final dataset due to low UMI per spot

H) DV200 = 54%, RIN: N/A, time since explantation: 4 years

I) DV200 = 17%, RIN: 2.50, time since explantation: 8 years

J) DV200 = 37%, RIN: 2.50, time since explantation: 5 years

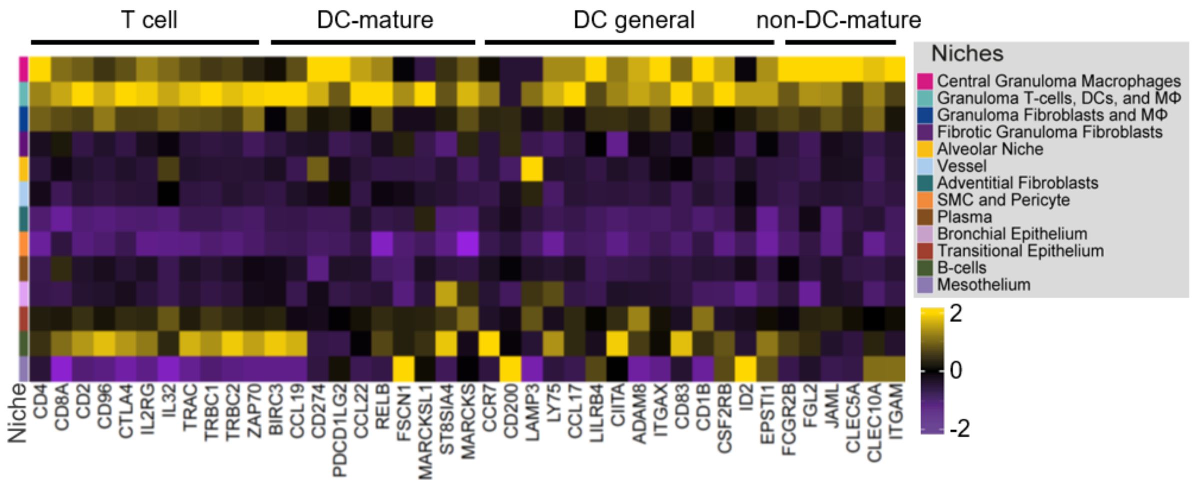


**Figure E2:** Heatmap displaying the expression of T cell and dendritic cell (DC) associated genes as $\log_2$(FC), grouped by their expression in T cells, mature DC, mature DC and other DC (DC general) and DC except mature-DC according to Schupp et. al (16).

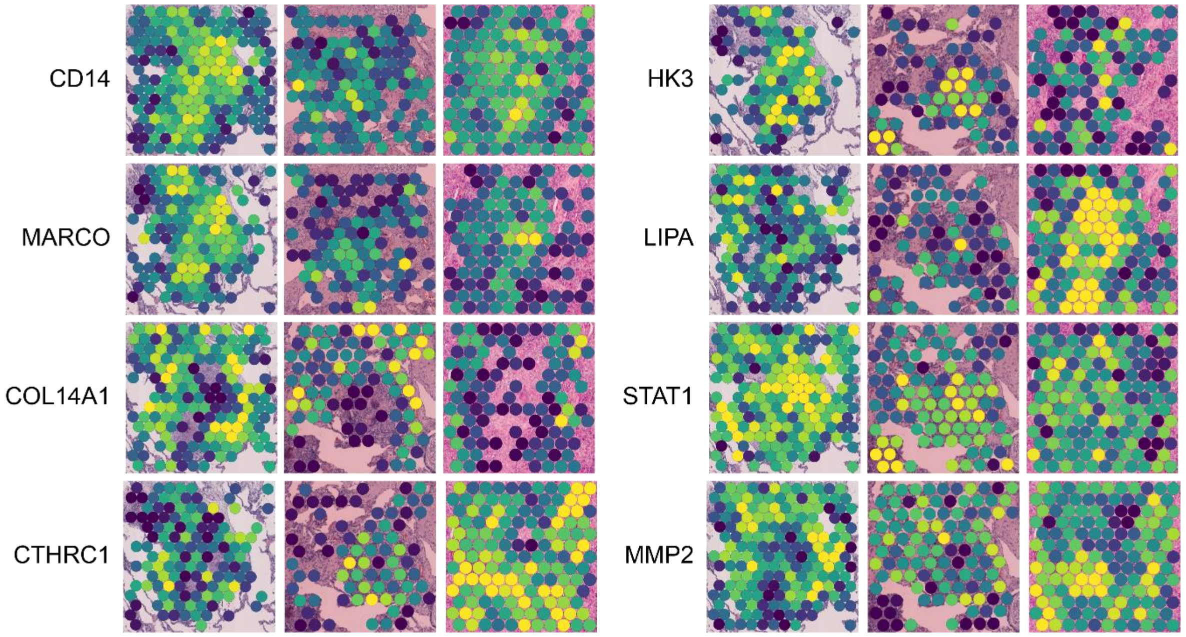


**Figure E3:** Spatial gene expression plot of macrophage- and fibroblast-related genes around a granuloma.

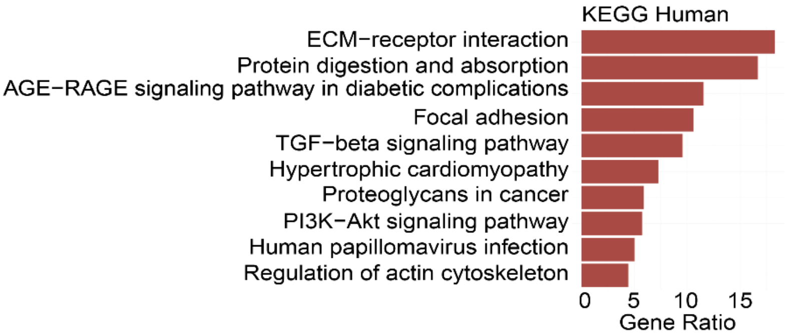


**Figure E4:** Pathway analysis of the top 200 genes expressed in the fibrotic granuloma fibroblast niche by $log_2$(FC). Reference database: KEGG.

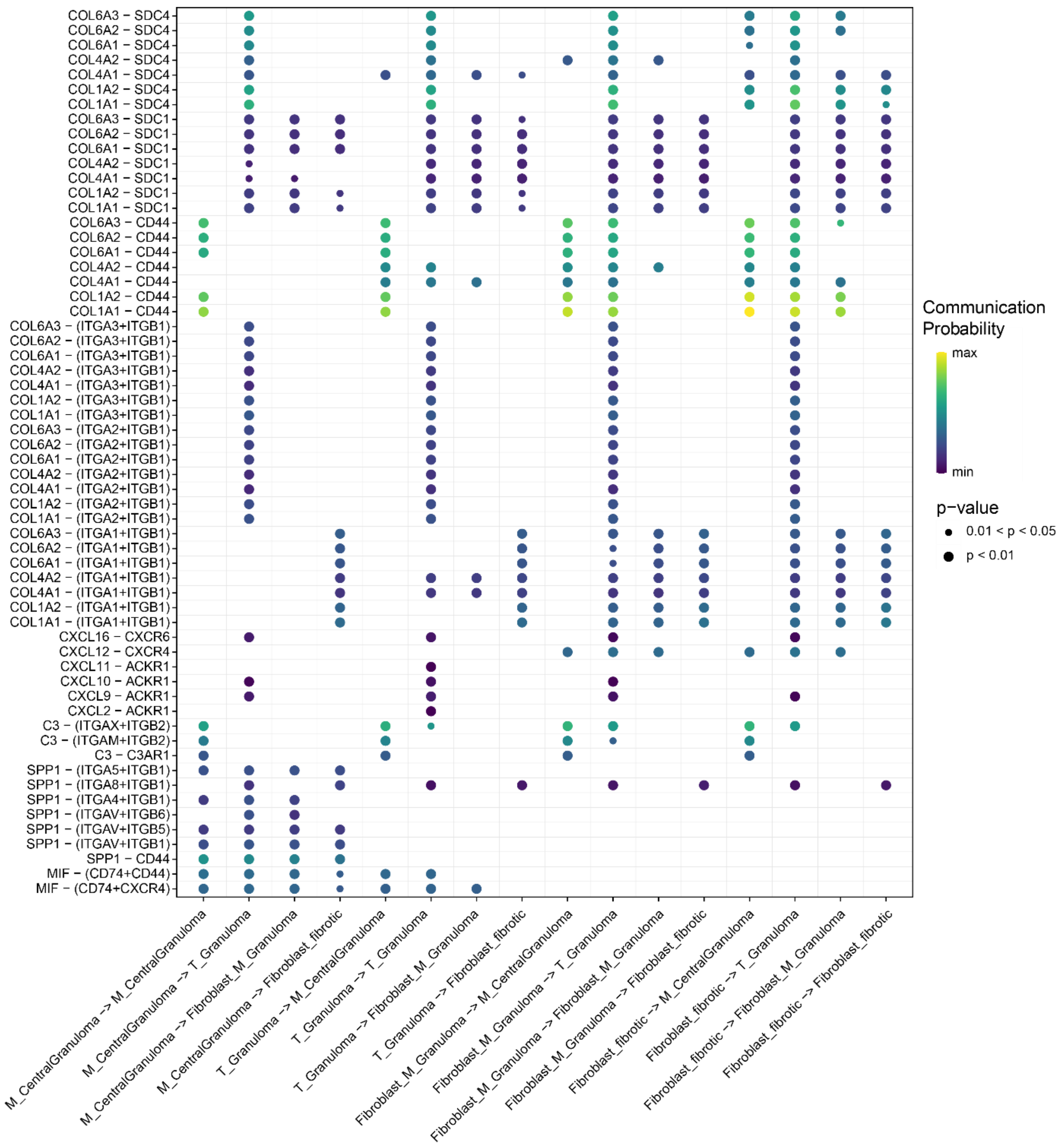


**Figure E5:** Bubble plot featuring the communication probability of each individual ligand-receptor interaction of any respective pathway featured in figure 6.

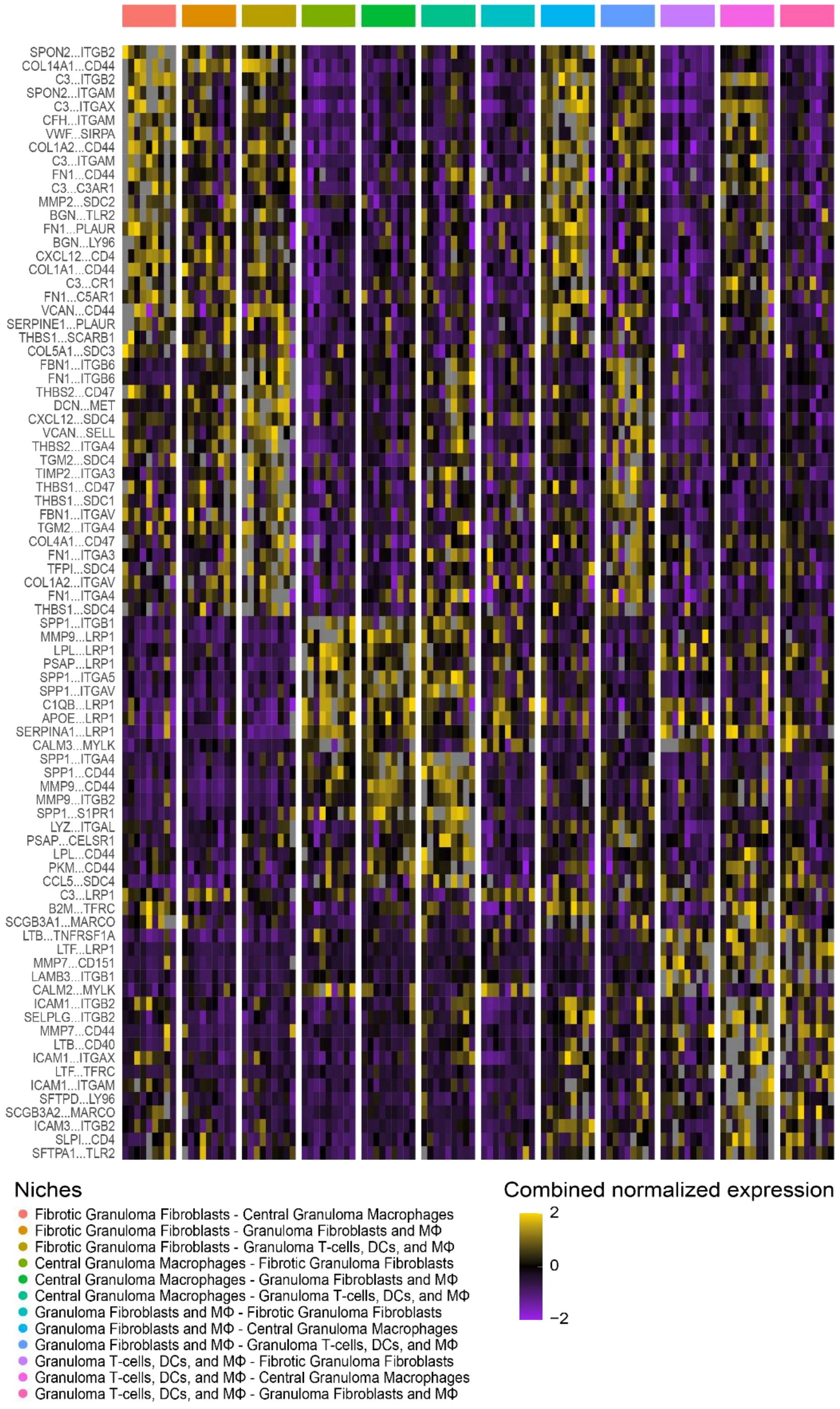


**Figure E6:** Combined normalized expression of ligand-receptor-pairs as average per subject of genes with a minimal differential gene expression of $log_2$(FC) = 0.3; Displayed are ligand-receptor-pairs having a granuloma-associated niches as a signal sender (ligand) with a different granuloma-associated niche as a receiver (receptor); nomenclature: ligand…receptor
